# Sulfated agarans from red seaweed *Gracilaria cornea* induce macrophages polarization to an antitumor M1 phenotype

**DOI:** 10.1101/2021.07.01.450714

**Authors:** Felipe Barros Teles, Alexia Nathália Brígido Assef, Renato Martins Andrade, Vitória Virgínia Magalhães Soares, Antônio Willame da Silva Alves, Roberto César Pereira Lima-Júnior, Norma Maria Barros Benevides, Diego Veras Wilke

## Abstract

Marine seaweeds are a rich source of sulfated polysaccharides with several biological activities, including antitumor effect. Some polysaccharides are also described to activate macrophages (Mϕs) to an antitumor M1-like phenotype. Here, we evaluated the capacity of sulfated galactans (SGs) extracts obtained from three seaweed species, *Gracilaria cornea* (Gc-E), *Gracilaria birdiae* (Gb-E), and *Solieria filiformis* (Sf-E), to activate the Mϕs antitumor M1 phenotype. The nitric oxide production, MHCII, and CD86 (M1 markers) were evaluated to screening the bioactive SGs profile on murine Mϕs (RAW 264.7 cells). The direct SGs antiproliferative effect was tested on melanoma B16-F10 cells. In another experimental setting, B16-F10 cells were incubated with a conditioned medium obtained from Mϕs exposed to SGs. The three SGs tested induced NO release. Sf-E directly inhibited B16-F10 cells proliferation compared with the saline group, but Gc-E and Gb-E failed to inhibit cell proliferation. Notably, a conditioned medium (CM) of Mϕs incubated with Gc-E and Sf-E, but not of Gb-E, inhibited the proliferation of B16-F10 cells. Gc-E also induced TNF-α release and increase of M1 markers such as iNOS, MHCII, and CD86. Therefore, Gc-E activates Mϕs to M1 phenotype, which in turn releases a factor that inhibits B16-F10 proliferation.

## Introduction

Depending on the stimulus, monocytes/macrophages (Mϕs) undergo functional reprogramming and may exhibit a spectrum of different phenotypes. Interferon-gamma (IFN-γ) alone or associated with microbial lipopolysaccharides (LPS) polarizes Mϕs for a classically activated M1 phenotype. On the other hand, IL-4 and IL-13 cytokines induce an alternatively activated M2 phenotype. Notably, M1-like phenotypes contribute to resistance against tumors, while the M2 Mϕs trigger protumor mechanisms (Mantovani et al., 2014).

Sulfated polysaccharides (SPs) are biological macromolecules with a high negative charge (Bedini et al., 2017; Ruocco et al., 2016). In red seaweeds SPs are represented by the sulfated galactans (SGs) (Cian et al., 2015). Depending on the seaweed species, galactans exhibit structural differences and are divided into agaranas or carrageenans (Usov, 2011). Previous studies have demonstrated *in vitro* and *in vivo* activities of SGs obtained from red seaweeds species, including agaran-type from *Gracilaria cornea* and *Gracilaria birdiae* (Usov, 2011) and carrageenan-type from *Solieria filiformis* (Robledo & Freire-Pleegrin, 2010). The biological activities include modulation of pain and inflammation (Coura et al., 2012), neuroprotective activity (Souza et al., 2016) anxiolytic (Monteiro et al., 2016), anticoagulant (Rodrigues et al., 2010), and antiviral effects (Morán-Santibañez et al., 2016). In this study we evaluated the activation and phenotype of murine macrophages RAW 264.7 incubated with SGs from red seaweeds *G. cornea*, *G. birdiae* and *S. filiformis*.

## Materials and methods

### Reagents

Dulbecco’s modified Eagle medium (DMEM), fetal bovine serum (FBS), antibiotics (penicillin and streptomycin), and trypsin-EDTA were purchase from Gibco BRL Co. (Grand Island, NY, USA). Lipopolysaccharide from *Escherichia coli* (LPS), acetic acid, triton X-100, propidium iodide, doxorubicin, sulfanilamide, and NED reagents were obtained from Sigma Aldrich (St. Louis, MO, USA). Sodium nitrite (NaNO_2_) was purchased from Dinâmica (Diadema, São Paulo, Brazil). TNF-α ELISA kit was purchased from R&D Systems (Minneapolis, MN, USA).

### Isolation and characterization of sulfated galactans

The seaweeds *G. birdiae* and *S. filiformis* were obtained during sampling procedures from an experimental culture located at 200 m from the coastal zone of Flecheiras Beach-Trairi, Ceará (03° 13‘06 “S; 039° 16’47” W). *Graciliria cornea* was collected in the intertidal zone at Flecheiras Beach. One voucher of each species (*G. cornea*, #34739; *G. birdiae*, #40781; and *S. filiformis*, #35682) was deposited in the Herbarium Prisco Bezerra, Department of Biological Sciences, Federal University of Ceará, Brazil. The seaweeds were washed with distilled water, separated from epiphytes or other fouling organisms, and stored at −20 °C until use. SGs derived from *G. cornea* (Gc-E) and *S. filiformis* (Sf-E) were obtained by protease digestion (60°C for six hours) in 100 mM sodium acetate buffer (pH 5,0) containing 5 mM EDTA and 5 mM cysteine, with some modifications as previously described (Coura et al., 2012a; Araújo et al., 2011). SGs derived from *G. birdiae* (Gb-E) was obtained by non-enzymatic aqueous extraction at 85 °C, according to Bezerra and Marinho-Soriano (Bezerra & Marinho-Soriano, 2010), with some modifications. The use of these algae species was registered on the National System of Management of Genetic Heritage and Associated Traditional Knowledge (SisGen) under the number A93CDDO.

### Cell lines, culture procedures and treatments

RAW 264.7, a murine macrophage cell line, and B16-F10, a murine metastatic melanoma cell line, were purchased from the Banco de Células do Rio de Janeiro (Rio de Janeiro, Brazil) and kept in the Laboratory of Marine Bioprospection and Biotechnology (LaBBMar) that has biosafety level 2 certified by National Technical Commission on Biosafety (CTNBio, Brazil) The cells were cultivated in DMEM containing 10% heat-inactivated FBS and 1% penicillin/streptomycin at 37°C in a 5% CO2 humidified atmosphere. The cells were regularly split to keep them into the exponential phase of growth. The Laboratory of Marine Bioprospection and Biotechnology at Drug Research and Development Center, Federal Univesrity of Ceará was granted by the National Technical Commission on Biosafety (CTNBio, Brazil) as a biosafety level 2 laboratory under the number 6.019/2018.

### NO analysis

RAW 264.7 cells were exposed for 24 h to Gc-E (10, 100, or 250 μg/mL, Sf-E (1, 10, or 100 μg/mL), Gb-E (10, 100, or 250 μg/mL), saline (as negative control) or LPS (100ng/mL), as positive control. The nitric oxide (NO) levels formed in the supernatant of macrophages RAW 264.7 were indirectly measured by the Griess’ test (Green et al., 1982). Briefly, 50 μl of cell culture supernatants were mixed with 50 μl of Griess reagent (0.1% [wt/vol] naphthyl ethylenediamine and 1% [wt/vol] sulfanilamide in a 5% [vol/vol] phosphoric acid) in a 96-well plate, incubated at room temperature for 10 min. The absorbance was then measured with a microplate reader at 570 nm. The absorbance values of the treated groups were interpolated from the linear regression performed with the data of the standard curve.

### Evaluation of antiproliferative effect in vitro

The antiproliferative assay of samples was performed by SRB assay (Skehan et al., 1990). SRB method is used to determine cell density based on the total intracellular protein content measurement and does not depend on cellular metabolism. B16-F10 cells were either incubated for 48h with saline, LPS (100 ng/mL), Gc-E, Sf-E, or Gb-E (1–250 μg/mL). In another experimental setting, RAW 264.7 cells were incubated for 24 h with the same compound concentrations mentioned above for obtaining the conditioned medium. B16-F10 cells were then incubated for 48h with conditioned medium obtained from saline-treated macrophages (CM-Sal), LPS-treated Mϕs (CM-LPS), or SG-treated macrophages (CM-Gc-E, CM-Sf-E, or CM-Gb-E). Therefore, we added 100 μL of RAW 264.7 CM to an equal volume of the B16-F10 supernatant (1:1 v/v), resulting in a dilution factor = 2.

### Cytokine assays

We measured TNF-α cytokine levels in supernatants of RAW 264.7 plated at 1.5 x 10^5^ cells/mL in 96 wells plate by ELISA, according to the manufacturers’ instructions. The absorbance values of the treated groups were interpolated from the linear regression performed with the data of the standard curve.

### Analysis of cell immunostaining by flow cytometry

We used murine FITC-conjugated anti-CD86 or murine PE-conjugated anti-MHCII antibodies from Thermo Fisher (Waltham, MA, USA) to stain the activation markers CD86 and MHCII, respectively. RAW 264.7 was incubated with the antibodies for 30 minutes at 4°C in FACS buffer (PBS supplemented with 4% FBS) and then washed with FACS buffer again. The cells were also fixed with paraformaldehyde 1% for 5 minutes and permeabilized with Triton 0.1% for 5 minutes for iNOS labeling. Then, cells were incubated with murine Alexa Fluor 488-conjugated antibody for 30 minutes at 4 °C in FACS buffer and washed with FACS buffer again. We used flow cytometry (model FACSVerse, BD Biosciences, San Jose, USA) for data acquisition and the FlowJo software (San Jose, CA, USA) for the analysis of the median fluorescence intensity values (MFI) of iNOS, MHCII, and CD86 expression, which were normalized by the mean of negative control MFI. The percentage of double-labeled cells was used to show the population of cells expressing MHCII and CD86 simultaneously.

### Statistical analysis

The data represents the mean ± SD of three technical and biological replicates. We used one-way analysis of variance (ANOVA) followed by Dunnett or Tukey’s multiple comparison tests to detect the statistical difference between the groups. *P* < 0.05 was considered statistically significant. Statistical analysis was performed using the GraphPad Prism 6.0 software (San Diego, CA, USA).

## Results and Discussion

### Sulfated galactans extracts stimulate RAW 264.7 to produce nitric oxide (NO)

All SGs extracts stimulated NO release by Mϕs (Fig. 1A). While Gc-E stimulation was observed at all concentrations, Sf-E predominantly induced NO release at lower concentrations (1 and 10 μg/mL) and Gb-E at the higher concentration (250 μg/mL) (Fig. 1A). NO is a crucial component of the host immune response against various pathogens such parasites, bacteria, and viruses (Bogdan et al., 2000; Xue et al., 2018). NO production is considered a classical M1 phenotype marker (Zhu et al., 2014). SGs from brown seaweed *Cystoseria indica* induced the release of NO from RAW 264.7 by activating the Toll-like 4 receptor (TLR4)/NF-κB and MAPKs signaling pathways (Bahramzadeh et al., 2019). Another sulfated polysaccharide from seaweed, *Nizamuddinia zanardinii*, similarly activated RAW 264.7 to secrete NO, TNF-α, and other cytokines (Tabarsa et al., 2020). Additional *in vitro* studies further reported that galactans also activate Mϕs to M1-like phenotype. The mechanisms seem to involve TLR4- and autophagy-driven macrophage pro-inflammatory phenotype, as demonstrated by the hetero-galactan from fungi *Flammulina velutipes* (Y. Meng et al., 2018). The consistency of such mechanism seems to be shared by other galactans, including those obtained from Panax ginseng flowers or fruiting bodies of *Cantharellus cibarius*, which activate RAW 264.7 Mϕs to increase phagocytosis and the release of NO, TNF-α, IL-6, IFN-γ, and IL-1β (Cui et al., 2020; Yang et al., 2019).

**Figure 1.**
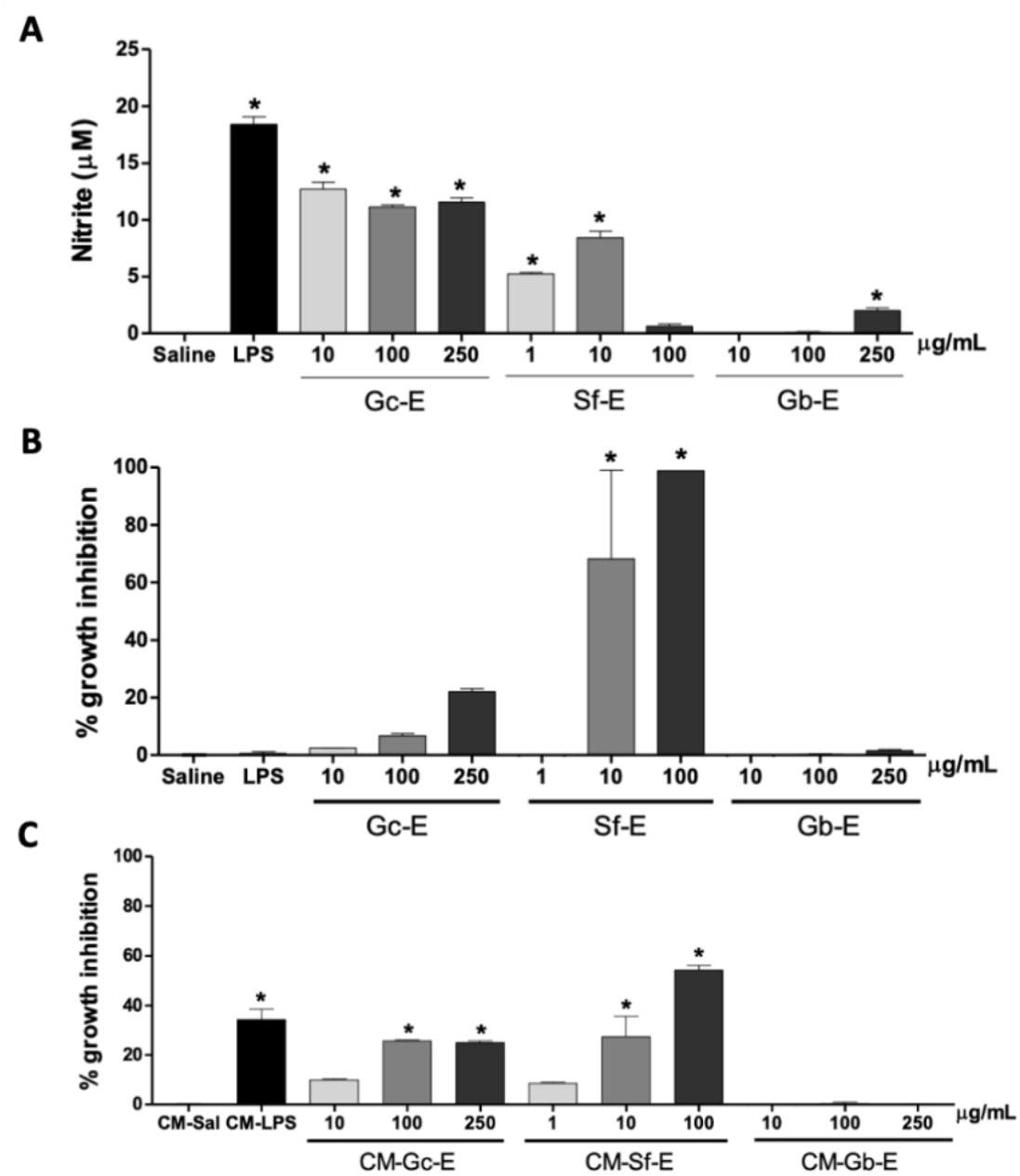
Bioactive profile of sulfated galactans extracts. **A**, Nitrite levels produced by macrophages (RAW 264.7 cell line) evaluated by Griess assay. RAW 264.7 cells were exposed to saline as a negative control, *E. coli* lipopolysaccharide (LPS) at 100 ng/mL as positive control and increasing concentrations of sulfated polysaccharides extracted from *Gracilaria cornea* (Gc-E), *Solieria filiformis* (Sf-E), or *Gracilaria birdiae* (Gb-E) for 24h. **B**, Antiproliferative effect of Gc-E, Sf-E, and Gb-E against the murine metastatic melanoma cell line (B16-F10) following a 48h exposure evaluated by SRB assay. **C**, Antiproliferative effect of macrophage-conditioned medium (CM) obtained after a 24-h RAW 264.7 cell incubation with Gc-E, Sf-E, or Gb-E. B16-F10 cells were exposed to CM for 48h. Differences between data normalized to percentage of cell growth inhibition (B and C) of negative control versus other groups were determined by the analysis of variance (ANOVA) followed by Dunnett’s post-test. *p<0.05.

### Antiproliferative effect of sulfated galactans extracts against B16-F10 cells

SGs are known for their antiproliferative effect on tumor cells (Khotimchenko et al., 2020). Accordingly, Sf-E demonstrated a marked antiproliferative effect on B16-F10 cells. Conversely, Gc-E and Gb-E failed to inhibit tumor cell growth (Fig. 1B). Remarkably, SGs of marine origin display low toxicity, which allows its widespread use as food compounds (Fedorov et al., 2013). Such findings contrast with the well-described *in vitro* antiproliferative activity of carrageenans. The iota-carrageenan extracted from red seaweed *Laurencia papillosa* inhibits cell proliferation of adenocarcinoma cell line by inducing apoptosis (Murad et al., 2015). The same is also described to the kappa(k)-carrageenan from red seaweed *Hypneia musciformis*, which reduces neuroblastoma cell-line proliferation (Souza et al., 2018). The mechanism might involve the phase-specific cell cycle arrest, as reported to kappa(κ) and lambda (λ)-carrageenans on HeLa cells (Prasedya et al., 2016).

### Conditioned medium obtained from RAW 264.7 cells stimulated with sulfated galactans extracts inhibits the proliferation of B16-F10 cells

The conditioned medium (CM) obtained from MΦs exposed to Gc-E (CM-Gc-E) or Sf-E (CM-Sf-E), but not CM-Gb-E, inhibited B16-F10 cell growth. The effective concentrations varied from 10 to 250 μg/mL depending on the extract (Fig. 1C). Engström and colleagues demonstrated that conditioned media of human macrophages (differentiated from THP-1 monocytes) polarized to an M1-phenotype by LPS and interferon γ inhibits the proliferation of colorectal adenocarcinoma cell lines (Engström et al., 2014). A comparable experimental approach also effectively inhibited the proliferation of a breast cancer cell line by macrophages derived from bone marrow (Rey-Giraud et al., 2012). Notably, as far as we know, this is the first report on the antitumor profile of MΦs activated by polysaccharides from natural sources. It is also worth to highlight two advantages conferred by this model used with Mϕs RAW 264.7 cell line instead of monocytes or bone marrow. Firstly, it was the overall shorter time to perform the experiments due to the lack of monocyte differentiation into macrophage step, and second, in the case of bone marrow as source of cells, it was the absence of animal ethical issues.

### Gc-E activates RAW 264.7 to produce TNF-α

Gc-E exhibited the most suitable bioactive profile among the tested SGs. It showed mild direct cytotoxicity against tumor cells, stimulated NO production by Mϕs, and stimulated the macrophages to release factors capable of inhibiting B16-F10 growth. Remarkably, Gc-E stimulated TNF-α production by RAW 264.7 at 100 and 250 μg/mL (**Fig. 2**). TNF-α is a cytokine produced by some immune cells, like Mϕs, during inflammation, infection, or injury and is also considered an M1 marker (Rock, C.S. & Lowry, S.F., 1994; Murray, P. et al., 2014). During the 70s, Carswell and collaborators demonstrated that TNF-α induces tumor regression (Carswell, E.A., 1975). To test B16-F10 responsiveness to TNF-α, we incubated these tumor cells with different concentrations of this cytokine (up to 1 ng/mL) to assess the inhibition of cell growth. Intriguingly, there was no cell growth inhibition at any of the concentrations tested (data not shown). Therefore, the antiproliferative factors released by Mϕs in the CM-Gc-E merit to be determined in a further study.

**Figure 2.**
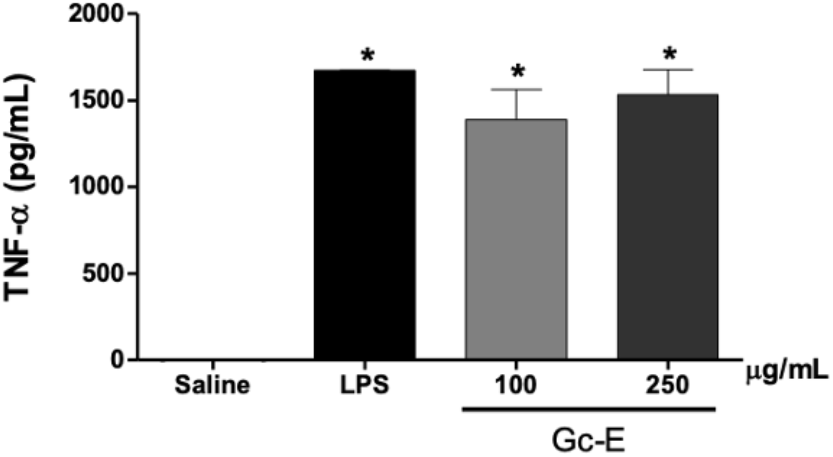
Sulfated agarans extract from *Gracilaria cornea* (Gc-E) induce TNF-α releasing by macrophages. RAW 264.7 cells were exposed to saline as a negative control, *E. coli* lipopolysaccharide (LPS, 100 ng/mL) as a positive control, or Gc-E (100 or 250 μg/mL). Differences between negative control versus the other groups were determined by the analysis of variance (ANOVA) followed by Dunnett’s post-test. *p<0.05.

### Gc-E increases the expression of macrophage activation markers

The inducible nitric oxide synthase (iNOS) is an isoform of the enzyme synthesizing NO using L-arginine as a precursor (Hevel et al., 1991; Marletta et al., 1988). The Griess assay detects nitrite levels as indirect evidence of NO production. Here, RAW 264.7 cells incubated with Gc-E (250 μg/mL) showed a five-fold increase in iNOS expression versus the saline group (Fig. 3A-B), which is in line with the nitrite levels (Fig. 1A), suggesting the involvement of NO in the Gc-E mechanism of action. Additionally, Gc-E at 250 μg/mL increased MHC class II and CD86 expressions by 3-fold and 1.5-fold, respectively, compared with the control group (Fig. 3 C-H). Macrophages and other immune cells express MHC class II molecules to present antigens for CD4+ effector T cells (Rock, K.L. et al., 2016), a process that involves CD86 co-stimulatory surface molecules (Porta, C. et al., 2015). Noteworthy, all these molecules are important markers of M1 macrophage activation (Xue et al., 2018).

**Figure. 3.**
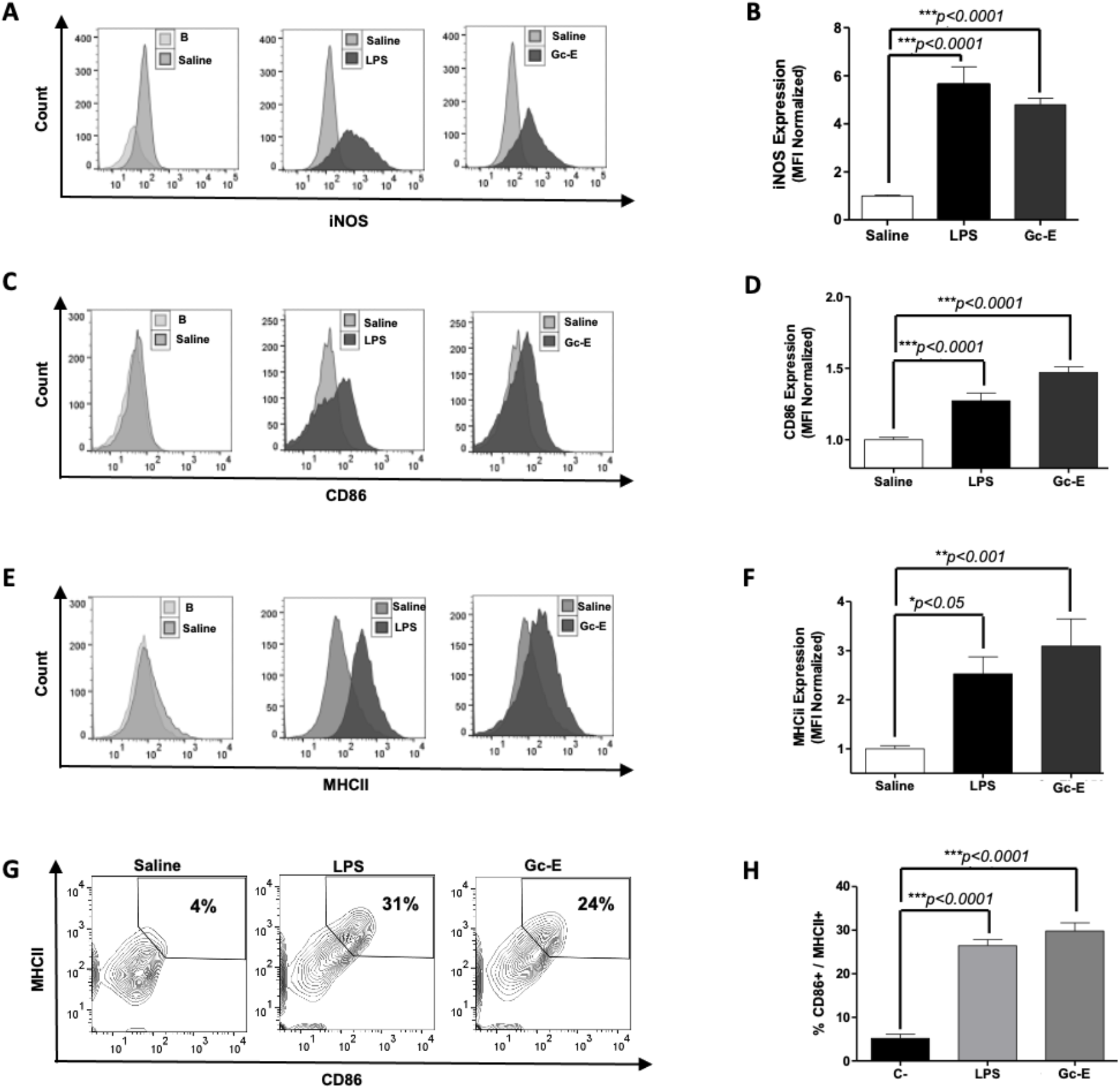
Sulfated agaran extract from *Gracilaria cornea* (Gc-E) induces a macrophage (RAW 264.7) M1 phenotype. Nitrite oxide production was evaluated by iNOS expression on RAW 264.7 cells by flow cytometry (A and B). Expression of CD86 and MHCII was evaluated on RAW 264.7 cells by flow cytometry (C–H). RAW 264.7 cells were incubated for 24 h with saline as a negative control, *E. coli* lipopolysaccharide (LPS, 100 ng/mL) as a positive control, and Gc-E at 250 μg/mL. Flow cytometry histograms are depicted on A, D, and E. Median fluorescence intensity (MFI) data are depicted on B, D, and F, as normalized values. Percentage of CD86+/MHCII+ cells are depicted on G and H. Ten thousand events, excluding debris and doublets, were acquired on flow cytometry experiments. Unstained cells were used as blank (B) as basal antibody background reference. The mean differences between the negative control versus other groups were compared by analysis of variance (ANOVA) followed by Dunnett’s post-test. *p<0.05. Results are representative of three independent experiments, each one performed in triplicate.

Gupta and cols. demonstrated that polysaccharides derived from *Tinospora cordifolia* induce the secretion of pro-inflammatory cytokines, such as TNF-α, IL-1β, IL-6, IL-12, and IFN-γ, by RAW 264.7. Moreover, NO levels are also enhanced along with the up-regulation of iNOS and surface MHC-II and CD-86 expression in murine Mϕs post-treatment with *T. cordifolia* polysaccharides (Gupta et al., 2017).

Meng and cols. showed that polysaccharide from a novel fungal strain isolated from the species *Cordyceps sinensis* promotes proliferation of RAW 264.7, increases phagocytic activity, releases NO, and increases the secretion of IL-6, Il-1α, IL-10, and TNF-α, and chemokines, such as MIP-1α and CXCL10 (IP-10). Both cytokines and chemokines are essential mediators released during the inflammatory processes. Eventually, the authors found enhanced expression of MHC II and CD86 in RAW 264.7 pre-treated with *C. sinensis* polysaccharides (L. Z. Meng et al., 2014). Remarkably, SPs from red seaweed *Champia feldmmanii* does not exhibit significant *in vitro* cytotoxicity but an *in vivo* antitumor effect. It might be explained by immunostimulating activity (Lins et al., 2009). Paclitaxel, a highly cytotoxic chemotherapeutic drug used to treat several cancers, also reduces tumor growth by reprogramming Mϕs to an M1 phenotype, a process mediated by Toll-like receptor-4 (Wanderley et al., 2018). Altogether, our results suggest that the polysaccharides from *G. cornea* are promising immunomodulatory agents. Additionally, Gc-E may exert *in vivo* antitumor inhibition by modulating the tumor microenvironment targeting Mϕs. Further studies are ongoing to address the immunostimulatory and *in vivo* antitumor effects of Gc-E.

## Conclusions

In summary, Gc-E induced macrophage activation to an M1 phenotype and depict antiproliferative *in vitro* effect on tumor cells. This antiproliferative effect of CM-GcE was not associated with direct cytotoxicity. Further studies are needed to understand the mechanisms involved in the activation of Mϕs by SGs and which factors are released by these cells in the conditioned medium to exert the GcE antiproliferative effect against B16-F10 cells.

## Acknowledgements

This work was supported by Coordenação de Aperfeiçoamento de Pessoal de Nível Superior - Brasil (CAPES) for scholarships and financial support (Ciências do Mar II CAPES finance code n° 23038.001422/2014-84) and National Institute of Science and Technology on Biodiversity and Natural Products (INCT BioNat-CNPq/FAPESP, Finance code n° 465637/2014-0). The authors also would like to thank the Multi-User Facility of Drug Research and Development Center of Federal University of Ceará for technical support.

